# Jointly embedding multiple single-cell omics measurements

**DOI:** 10.1101/644310

**Authors:** Jie Liu, Yuanhao Huang, Ritambhara Singh, Jean-Philippe Vert, William Stafford Noble

## Abstract

Many single-cell sequencing technologies are now available, but it is still difficult to apply multiple sequencing technologies to the same single cell. In this paper, we propose an unsupervised manifold alignment algorithm, MMD-MA, for integrating multiple measurements carried out on disjoint aliquots of a given population of cells. Effectively, MMD-MA performs an *in silico* co-assay by embedding cells measured in different ways into a learned latent space. In the MMD-MA algorithm, single-cell data points from multiple domains are aligned by optimizing an objective function with three components: (1) a maximum mean discrepancy (MMD) term to encourage the differently measured points to have similar distributions in the latent space, (2) a distortion term to preserve the structure of the data between the input space and the latent space, and (3) a penalty term to avoid collapse to a trivial solution. Notably, MMD-MA does not require any correspondence information across data modalities, either between the cells or between the features. Furthermore, MMD-MA’s weak distributional requirements for the domains to be aligned allow the algorithm to integrate heterogeneous types of single cell measures, such as gene expression, DNA accessibility, chromatin organization, methylation, and imaging data. We demonstrate the utility of MMD-MA in simulation experiments and using a real data set involving single-cell gene expression and methylation data.

## 1 Introduction

Next-generation sequencing has enabled high-throughput interrogation of many different physical properties of the genome, including the primary DNA sequence but also the expression of messenger RNAs, localized binding of specific factors, histone modifications, nucleosome occupancy, chromatin accessibility, etc. Most of these sequencing assays have been performed on populations of cells. However, such bulk measurements do not allow for easy characterization of systematic or stochastic variations in physical properties of the genome among cells within a given population. Over the past several years, a variety of genomic assays have been modified to allow characterization of single cells. These modifications sometimes involve physically segregating individual cells prior to sequencing, or alternatively involve successive rounds of DNA bar-coding to identify reads derived from single cells. Examples of single-cell genomics assays include single-cell RNA-seq (scRNA-seq) for gene expression [14], single-cell ATAC-seq (scATAC-seq) for chromatin accessibility [3], single-cell Hi-C (scHi-C) for 3D genome organization [10] and single-cell methylation analysis (scMethyl-seq) [13]. In each case, the result is a data set that, compared to a standard, bulk genomic assay, has an additional dimension corresponding to the cells in the sample population.

Single-cell measurements are valuable because they permit a view of the cell-to-cell variation of a given type of physical measurement of the genome. However, such measurements would be even more valuable if multiple different measurements could be obtained from the same individual cell. Such co-assays are feasible, albeit challenging and lower throughput, for pairs of assays, such as scRNA-seq and scATAC-seq [4] or scRNA-seq and scMethyl-seq [2], that measure orthogonal physical properties. However, other pairs of single-cell assays, such as scATAC-seq and scHi-C cannot be paired even in principle, because each assay operates on (and cleaves) the genomic DNA.

In this paper, we propose a manifold alignment algorithm based on the maximum mean discrepancy (MMD) measure, called MMD-MA, which can integrate different types of single-cell measurements. Our MMD-MA algorithm assumes that the cells are drawn from the same initial population—e.g., cells of the same type or a distribution of cell types from the same experimental conditions—but the algorithm does not require any correspondence information either among samples or among the features in different domains. The algorithm makes no parametric assumptions about the forms of the distributions underlying the various measurements. The only assumption is that the distributions share a latent structure with sufficient variability that the MMD term in the optimization can align the distributions. For example, if both underlying distributions are simple isotropic Gaussian distributions, then it will not be possible to reconstruct the relative orientation of the alignment. In practice, visualization of many different single-cell data sets using dimensionality reduction methods such as PCA, t-SNE or UMAP suggest that they commonly exhibit complex structure that, we hypothesize, should allow for alignment across data modalities. In particular, MMD-MA can be applied to many heterogeneous types of single cell measures, including gene expression, DNA accessibility, chromatin organization, methylation, and imaging data. Thus, the algorithm allows us, in principle, to obtain the insights offered by a single-cell co-assay by computationally integrating two or more separate sets of single-cell measurements derived from the same or similar populations of cells. We demonstrate the performance of the algorithm on three simulated data set as well as one real data set consisting of gene expression and methylation profiles of single cells.

## 2 Methods

### 2.1 The unsupervised manifold alignment problem

Our goal is to automatically discover a manifold structure that is shared among two or more sets of points that have been measured in different ways, i.e., which mathematically live in different spaces. For simplicity, we describe the case where we have just two types of measurements, though the approach generalizes to any number of input domains. Let the two sets of points be 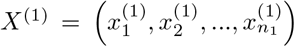 from 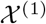 and 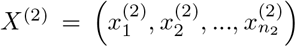 from 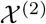. The numbers of data points in the two domains are *n*_1_ and *n*_2_, respectively. We do not require any correspondence information regarding the measurements across different domains or regarding the samples from different domains. Instead, we assume that both sets of points share a manifold structure, which we aim to discover in an unsupervised fashion.

To ensure the generality of our approach, we frame the optimization using kernels. Hence, we assume that we have a way of calculating similarities between pairs of entities from the same domain, using positive definite kernel functions 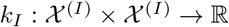 for *I* = 1, 2. The resulting kernel Gram matrices are denoted by 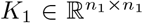 and 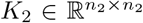, where 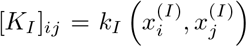 for *I* = 1, 2 and 1 ≤ *i, j* ≤ *n*_*I*_. As long as both kernel functions are positive definite, then we are guaranteed that each kernel corresponds to the scalar product operation in some induced feature space, and that there exists a space of functions 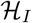, called a reproducing kernel Hilbert space (RKHS), mapping 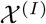 to ℝ endowed with a Hilbert space structure. For example, if the input space 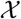 is a vector space and we take the linear kernel *k*(*x, x′*) = *x*^┬^*x′*, then the RKHS is made of linear functions of the form *f* (*x*) = *w*^┬^*x*, endowed with the norm ||*f*|| = ||*w*||. If we take a nonlinear kernel such as the Gaussian RBF kernel *k*(*x, x′*) = exp(−||*x* − *x′*||^2^/(2*σ*^2^)) with bandwidth *σ* > 0, then the RKHS contains nonlinear functions. The use of kernels allows the MMD-MA algorithm to operate in principal on any type of entity—vector, graph, string, etc.— for which a kernel function can be defined.

In order to find a shared structure between the points in 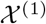 and 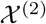, we propose to learn two mappings 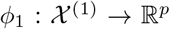 and 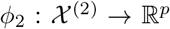, so that input data in different spaces are all mapped to the same *p*-dimensional space ℝ^*p*^ and can be compared in that space. For each *I* = 1, 2, we consider each coordinate of *φ*_*I*_ in the RKHS 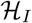 of the corresponding kernel *k*_*I*_, i.e., 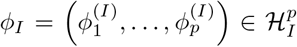. We then consider each function 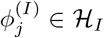 of the form 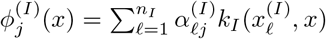, for any 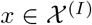. This parametrization of 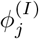 in terms of 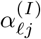’s always exists by the representer theorem, provided we regularize the optimization problem with the RKHS norm of 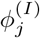, as we explain below. Now, if we denote by *α*_*i*_ the *n*_*i*_ × *p* matrix with entries 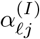, then *K*_*I*_*α*_*I*_ is the *n*_*I*_ × *p* matrix where the *j*-th row (for *j* = 1, …, *n*_*I*_) is the *p*-dimensional image of 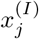 by the mapping *φ*_*I*_. In addition, 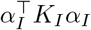 is the *p* × *p* matrix of inner products in the RKHS of the *p* functions 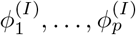, which is for example equal to the *p* × *p* identity matrix *I*_*p*_ when *φ*_*I*_ is a projection onto a subspace of dimension *p* in the RKHS. In order to define MMD-MA, we now discuss the criteria to optimize for *α*_1_ and *α*_2_ in order to discover shared structures between the two views.

### 2.2 Characterizing the distribution distance in the shared space

Although we do not assume we know the individual correspondence between points in the two domains, or even that such a 1-to-1 mapping exists, we do assume that the two distributions of points are similar in the shared space. Thus, the optimal mapping matrices *α*_1_ and *α*_2_ should make the two mapped sets of points in the shared space, namely *K*_1_*α*_1_ and *K*_2_*α*_2_, as similar as possible. To specify the distance between the two mapped manifolds in the shared space, we use an MMD term MMD(*K*_1_*α*_1_*, K*_2_*α*_2_)^2^, which is a general, differentiable measure of how similar two clouds of points are [7]. Formally, MMD is defined through a positive definite kernel *K*_*M*_ over ℝ^*p*^ through the formula

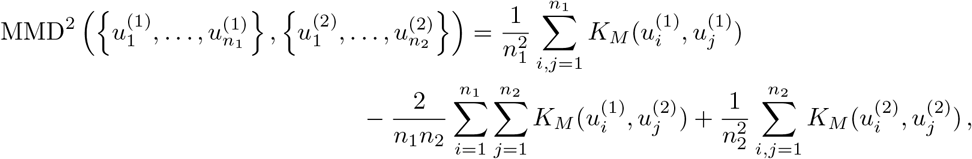

where we denote 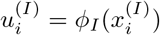 to simplify notation. In this work, we use a Gaussian RBF kernel for *K*_*M*_, where the bandwidth parameter *σ* is a user-specified parameter.

Chwialkowski et al. [5] propose two fast methods (with complexity linear in *n*_1_ + *n*_2_) to estimate MMD^2^, both of which are differentiable with respect to the positions of the points. Because the MMD is small when the distributions are similar, MMD-MA aims to minimize MMD^2^ with respect to the embeddings.

### 2.3 The MMD-MA algorithm

Unfortunately, simply minimizing MMD between the two kernels is insufficient. Most notably, we need to ensure that the relationships among data points in the input space is preserved to some extent in the feature space; otherwise, the method may learn very complicated mappings that completely modify the relative positions of cells in order to have them aligned between the two views. For that purpose we introduce a distortion term 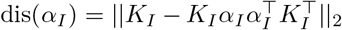, which quantifies how the matrix of inner products between points in the original space (quantified by the kernel matrix *K*_*i*_) differs from the matrix of inner products after mapping in the *p*-dimensional space. Penalizing dis(*α*_*I*_) ensures that the distortion between the data in the original space and the data mapped to the low-dimensional space should be small. In addition, we may wish to ensure that the mappings to ℝ^*p*^ are (almost) projections from the high-dimensional RKHS, which we obtain by adding a penalty term 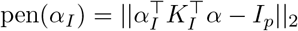.

Thus, MMD-MA optimizes, with respect to *α*_1_ and *α*_2_, the following objective function:

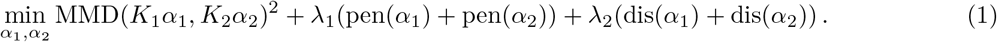

wher

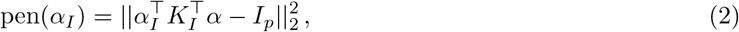

and

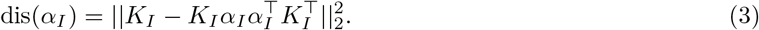

### 2.4 Solving the optimization problem

To find a stationary point of (1) we use a simple gradient descent scheme, and use Adam [9] to adjust the learning rate. Since the optimization problem is not convex, only a local minimum can be expected. Therefore, for a given data set we run the optimization procedure with 100 different random seed values and keep the solution which provides the lowest objective function value.

In practice, solving the optimization requires specifying several hyperparameters. These include the dimensionality *p* of the shared space, any parameters of the kernel functions *K*_1_, *K*_2_ and *K*_*M*_, and the tradeoff parameters *λ*_1_ and *λ*_2_. In this work, we assume that *p* and the kernel parameters are user-specified, and we investigate the performance of the algorithm as we vary *λ*_1_ and *λ*_2_.

## 3 Related work

We are aware of three other methods that address the unsupervised manifold alignment problem, which we briefly review here.

### 3.1 The joint Laplacian manifold alignment (JLMA) algorithm

The joint Laplacian manifold alignment (JLMA) algorithm [15] performs manifold alignment by constructing a joint Laplacian across multiple domains and then performing eigenvalue decomposition to find the optimal solution. The joint Laplacian formulation can also be interpreted as preserving similarities within each view and correspondence information about instances across views, which is captured by the joint Laplacian matrix. The loss function is *C*(*F*) = Σ_*i,j*_||**F**(*i,.*) − **F**(*j,.*)||^2^**W**(*i, j*), where the summation is over all pairs of instances from all views. **F** is the unified representation of all instances, and the output of the algorithm is the joint adjacency matrix **W**. To avoid trivial solutions (i.e., mapping all instances to zero), JLMA includes a constraint **F**^′^**DF** = *I* where *I* is an identity matrix. Let **F** = [*f*_1_, *f*_2_, …, *f*_*d*_]. The optimization problem then becomes

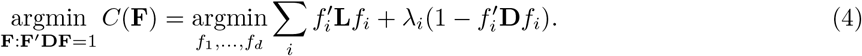

The optimal solution is the *d* smallest nonzero eigenvectors from the generalized eigen decomposition problem.

JLMA can be used in an unsupervised or supervised fashion. In supervised mode, the Laplacian *L* is given as input. In the unsupervised setting, the key step is to construct the cross-domain Laplacian submatrix of **L**. Wang *et al.* use *k*-NN graphs to characterize local geometry and use the minimum distances from scaled permutated *k*-NN graphs to construct a cross-view Laplacian submatrix of **L**. Thereafter, the rest of the algorithm is the same as the standard JLMA algorithm. Unfortunately, the computational cost of this initial step is quite high, even for small *k* values. To deal with this problem, Pe *et al.* use a B-spline curve to fit the local geometry and calculate cross-view matching scores from the curves [11]. Thus, in both cases, unsupervised manifold alignment is done via two steps: computing a cross-domain matching score, and identifying the correspondence via Equation 4.

### 3.2 The generalized unsupervised manifold alignment (GUMA) algorithm

The generalized unsupervised manifold alignment (GUMA) algorithm [6] is another method that does not require any correspondence information a priori. The approach assumes that instances in the two domains (e.g., in our case, two cells measured using different techniques) can be matched to one another in a one-to-one fashion. In particular, the algorithm formulates an optimization problem whose objective function includes a geometry matching term *E*_*s*_ across different domains, a feature matching term *E*_*f*_, and a geometry preserving term *E*_*p*_, subject to a 0-1 correspondence matrix **F** and feature projections **P**_*i*_ for each domain (i.e. domain *i*). The optimization is performed using alternating minimization, alternating between optimizing **F** and **P**_*i*_ with the other fixed. The algorithm outputs both the instance correspondence between the domains and the feature mapping functions between the domains.

### 3.3 The manifold alignment generalized adversarial network (MAGAN) algorithm

MAGAN [1] consists of two GANs that learn reciprocal mappings between two domains; i.e., GAN1 learns the mapping from domain 1 to domain 2, and GAN2 learns the mapping from domain 2 to domain 1. Each GAN’s generator takes input in one domain and outputs in the other domain, with the hope that the discriminator in the other domain cannot distinguish the fake samples from true samples. The loss function of the generators consists of three terms. The reconstruction loss term *L*_*r*_ captures the difference between a sample and itself after being mapped to the different domain and then mapped back to the original domain. The discriminator loss term *L*_*d*_ makes sure that the mapped sample in the other domain has a high likelihood to fool the discriminator in that domain. And the correspondence loss *L*_*c*_ forces the learned mapping to agree with some prior correspondence, either “unsupervised correspondence” (e.g., some variables are shared between two domains) or “semi-supervised correspondence” (e.g., some labeled pairs cross two domains). The paper empirically demonstrates that the inclusion of correspondence information greatly improves the performance of the manifold alignment.

### 3.4 Comparison of these three algorithms with our algorithm

The MMD term in our formulation only ensures that the two distributions agree globally in the latent space, whereas both JLMA and GUMA have a term that ensures, for each instance, that the local geometry is preserved between domains. This is the difference between *manifold superimposing* [18, 17, 8] and *manifold alignment* discussed in the MAGAN paper [1]. Furthermore, GUMA’s assumption that individual cells can be matched 1-to-1 between the two input domains is not generally true, most obviously when *n*_1_ ≠ *n*_2_. MAGAN [1] itself does not include a component for identifying a correspondence in an unsupervised fashion, and empirical results from the MAGAN paper suggest that the algorithm may not be useful if there is no known correspondence information between the two domains. When more than three domains must be aligned, the JLMA, GUMA, and our MMD-MA algorithms can be easily extended, whereas the formulation of MAGAN makes such an extension difficult.

## 4. Results

### 4.1 Three simulations

To validate the performance of MMD-MA, we generated three simulated data sets, each from a different *d*-dimensional manifold. The first manifold exhibits a branching structure in two-dimensional space (i.e., *d* = 2) to mimic a branching differentiation situation (first column of Figure 1). The second manifold structure is a nonlinear mapping of the first structure. The branching structure is mapped onto a Swiss roll manifold (second column of Figure 1). Samples in the first domain are mapped from the 2D space of simulation 1 into 3D space such that the three dimensions are [*x*_1_*cos*(3*x*_1_), *x*_2_, *x*_1_*sin*(3*x*_1_)], while samples in the second domain are mapped into 3D space by [*x*_1_*sin*(2*x*_1_), *x*_2_, *x*_1_*cos*(2*x*_1_)]. The third manifold is a circular frustum in three-dimensional space (i.e., *d* = 3), which aims to mimic the cell cycle superimposed on a linear differentiation process (third column of Figure 1). From each of these *d*-dimensional manifolds, we simulated *n* = 300 data points, and we refer to the corresponding *n* × *d* data matrix as *Z*.

**Figure 1:**
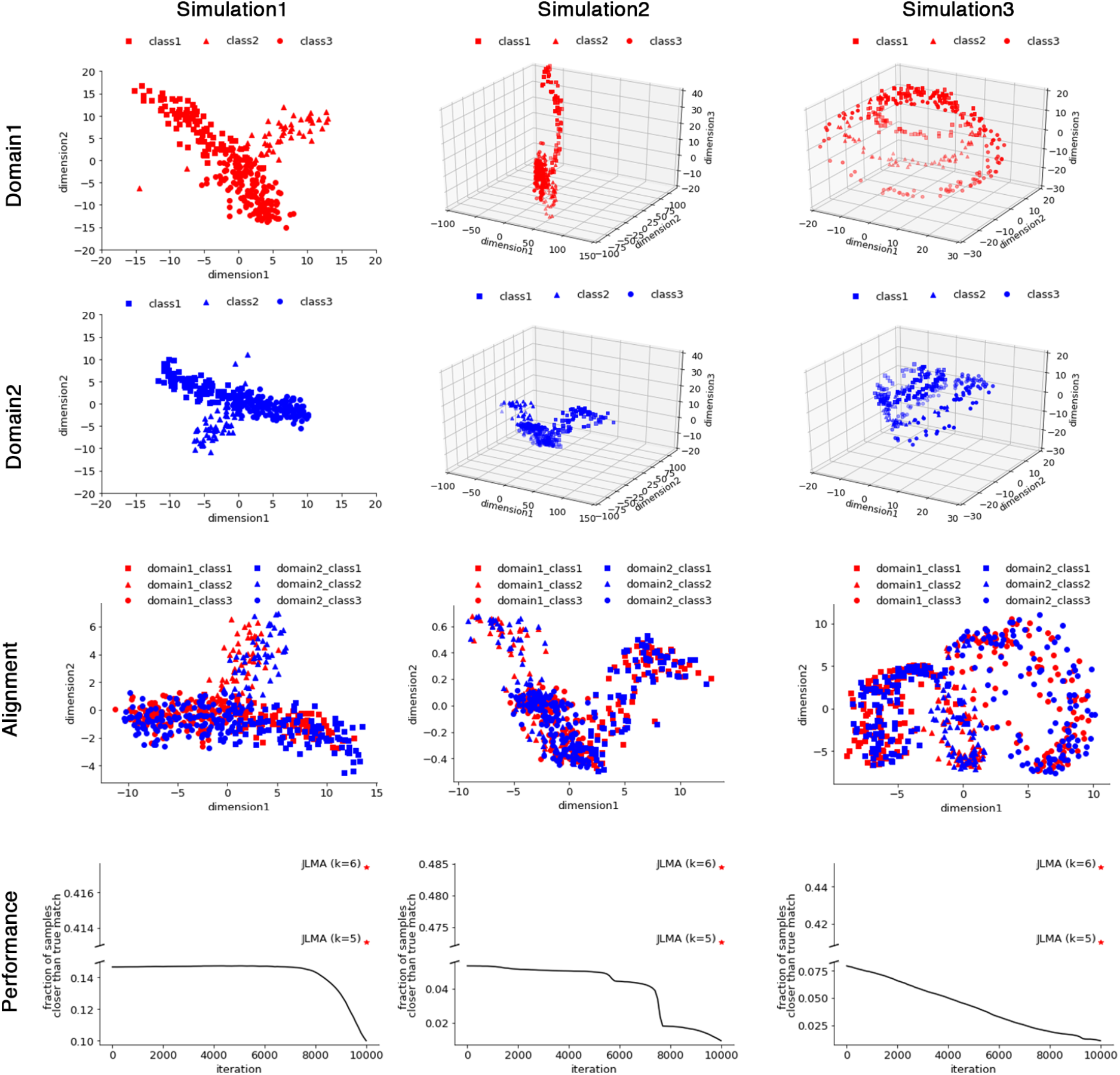
Three simulation experiments. The first two rows show the MDS projection of the data points in domain 1 and domain 2, separately. The third row shows the projection of the data points in the shared embedding space. The last row plots the fraction of samples closer than the true match as MMD-MA iterates. Points are included from JLMA when *k*=5 and *k*=6.

For each manifold, we assume that we have two domains, and we generate a *d × p*_1_ mapping matrix *T*_1_ and a *d* × *p*_2_ mapping matrix *T*_2_. Each element in the two mapping matrices is sampled from a standard Gaussian distribution. We set the observed data matrix from domain 1 to be *ZT*_1_ and the observed data matrix from domain 2 to be *ZT*_2_. For example, we set *p*_1_ = 1000 and *p*_2_ = 2000. We also add Gaussian noise (*σ* = 0.05) to each element of the covariates. For all of the simulations, we set the kernel matrices *K*_1_ and *K*_2_ to be the inner product of the *z*—normalized observed data matrices. For each simulation, we intentionally mis-specify, as input to MMD-MA, the dimensionality of the latent space as *p* = 5, to simulate the scenario in which the true latent dimensionality is unknown a priori.

For all three numerical simulations, we plot the data points in the projected space, namely 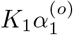 and 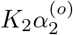 (Figure 1). In all three cases, the two domains appear to be aligned correctly in the latent space. To quantify this alignment, we use the known correspondence between points in the two domains as follows. For each point *x* in one domain, we identify its (true) nearest neighbor in the other domain. We then rank all data points in the learned latent space by their distance from *x*, and we compute the fraction of points that are closer than the true nearest neighbor. Averaging this fraction across all data points in both domains yields the “average fraction of samples closer than the true match,” where perfect recovery of the true manifold structure yields values close to zero. In all three simulations, the observed fraction of samples closer than the true match decreases monotonically and approaches zero as the MMD-MA algorithm iterates.

Finally, we attempted to compare the performance of other algorithms on the same simulated data sets. Unfortunately, we had difficulty running the GUMA algorithm using the implementation shared by the authors; hence, we leave GUMA out of our comparison. Similarly, we could not include the MAGAN algorithm because it requires some initial correspondence information, which we are assuming is not available. Consequently, we only compare to the JLMA algorithm as a baseline, using *k* = 5 (the default value) and *k* = 6. We find that, in all three simulations, our MMD-MA algorithm outperforms the baseline JLMA (bottom row of Figure 1).

The running time of MMD-MA is much lower than JLMA using either *k* = 5 or *k* = 6. Timings on an Intel Xeon Gold 6136 CPU at 3.00GHz (Table 1) show that MMD-MA runs under one minute, considerably faster than JLMA.

**Table 1:** Running time of MMD-MA and JLMA. Times are provided for the three simulation experiments and the real single cell application.

|  | Sim 1 | Sim 2 | Sim 3 | Methyl-Expr |
| --- | --- | --- | --- | --- |
| MMD-MA | 0:53 | 0:58 | 0:59 | 0:10 |
| JLMA, $k = 5$ | 4:06 | 4:07 | 4:20 | 1:27 |
| JLMA, $k = 6$ | 26:11 | 25:56 | 32:16 | 2:16 |

### 4.2 Real world application results

In a recent study, gene expression levels and methylation rates were profiled jointly in 61 single cells [2]. We use this co-assay data to validate our method by hiding the correspondence between genes from MMD-MA and then measuring how well the correspondence is recovered. Prior to analysis, we remove the genes where any of the cells have a missing value for either the methylation rate or gene expression. This step leaves, for the 61 cells, 2486 genes with both methylation rate and gene expression measured. We regard gene expression as domain 1 and methylation rate as domain 2. We pretend that we do not know the correspondence information, run our MMD-MA algorithm, and see how well our algorithm can align the two manifolds and recover the cell correspondence. For calculating the similarity kernel matrices *K*_1_ and *K*_2_, we first perform *z*-score normalization on the gene expression levels and the methylation rates and then calculate the inner product for the elements in the cell-by-cell similarity matrices. As in the simulations, we embed the two domains into a latent space of dimensionality *p* = 5.

We first plot the Principal Component Analysis (PCA) projection of the single cells based on their gene expression levels and their methylation rates separately (Figure 2A). In this plot, when we connect the two dots corresponding to the same cell, we observe that each cell tends to be projected to two different locations in the latent space. Accordingly, the average fraction of data points closer than the true match is 0.49. We then run MMD-MA algorithm on this dataset and plot the PCA projection of the 61 single cells in terms of gene expression and methylation rate in the shared space recovered by MMD-MA (Figure 2B). In the shared space projection, we connect the two embeddings from different perspectives, and we observe that the cells are embedded well in the shared space. Next, we calculate the fraction of samples closer to each cell that its true match in the shared space of dimensionality *p* = 5. This fraction decreases as MMD-MA iterates, reaching 0.024 in the end, and the trend is consistent across different learning rates of the optimization (Figure 2C). An alternative visualization of the per-cell fractions before and after optimization (Figure 2D) further illustrates that the MMD-MA algorithm successfully maps *>*50% of the cells closest to their true neighbor in the other domain.

**Figure 2:**
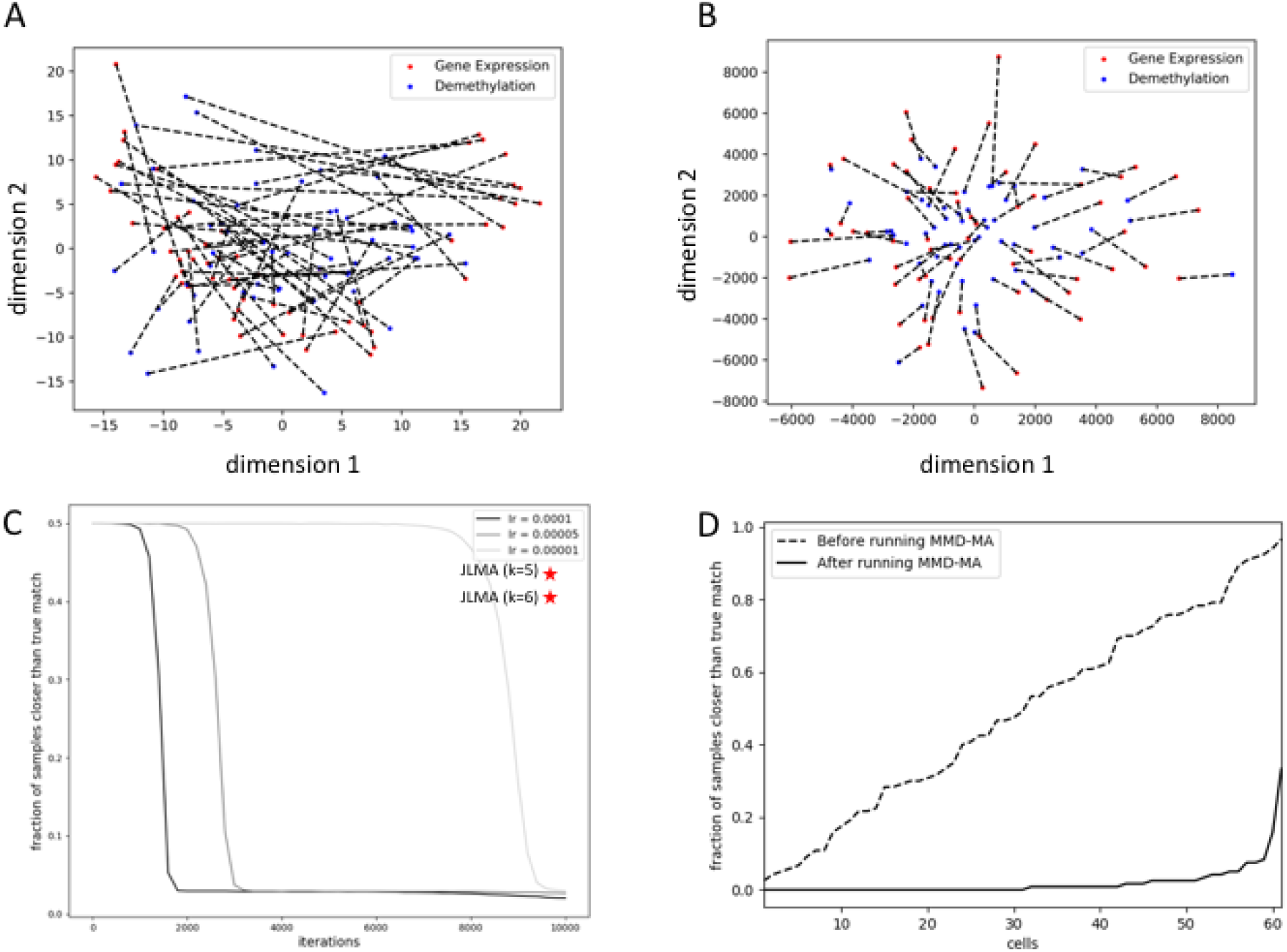
Results from real world single cell applications. (A) PCA projection of single cells based on their gene expression levels and their methylation rates separately, with dotted lines connecting the same cell. (B) Projection of the single cells in the shared space from the MMD-MA algorithm, with dotted lines connecting the same cell. (C) The average fraction of samples closer than the true match decreases as MMD-MA iterates. This result is consistent across different learning rates of the optimization. (D) The fraction of samples closer to each cell than its true match is plotted before and after MMD-MA, with the 61 cells in sorted order along the x-axis. For each cell, the average is computed separately for each domain, and then the two values are averaged together. The fraction is high and close to uniformly distributed before running the MMD-MA algorithm and reduces considerably as the algorithm learns the aligned shared space.

### 4.3 MMD-MA’s performance is robust to variations in hyperparameters

Running the MMD-MA algorithm requires specifying several hyperparameters. We investigated the robustness of the learned embedding relative to variations in these hyperparameters.

As noted previously, in all of our studies the dimensionality *p* of the latent space has been set to 5 even though the correct number should be *p* = 2 in the first two simulations, *p* = 3 in the third simulation, and is unknown for the Methyl-Expr data set. We observe that MMD-MA algorithm can still align the two manifolds even when the dimensionality parameter *p* is misspecified.

The trade-off parameters *λ*_1_ and *λ*_2_ determine how much the three terms contribute to the overall objective function. In this work, we set these trade-off parameters by monitoring whether the three terms have comparable magnitudes or whether one particular term dominates in the converged solution. We tested eight combinations of these trade-off parameters for each data set (Supplementary Table S1). In each case, we observe that the performance of MMD-MA is almost the same with different choices of trade-off parameters, although some trade-off parameters may lead to a different convergence path (Supplementary Figure S1 A–D).

The bandwidth parameter *σ* associated with the Gaussian kernel *K*_*M*_ in the MMD term determines how much each data point contributes to its neighborhood in the calculation of the MMD. The *σ* values used in our experiments are shown in Table 2. We also tested different values of *σ* and observed that the performance of MMD-MA is quite invariant to them (Supplementary Figure S1 E–H).

**Table 2:** Properties and hyperparameters of the experiments.

|  | Sim 1 | Sim 2 | Sim 3 | Methyl-Expr |
| --- | --- | --- | --- | --- |
| number of Samples | 300 | 300 | 300 | 61 |
| dimension (Domain 1) | 1000 | 1000 | 1000 | 2486 |
| dimension (Domain 2) | 2000 | 2000 | 2000 | 2486 |
| $\lambda_1$ | 1e-6 | 1e-9 | 1e-5 | 1e-2 |
| $\lambda_2$ | 1e-2 | 1e-7 | 1e-6 | 1e-6 |
| $\sigma$ | 0.5 | 0.1 | 1.2 | 10000 |

## 5 Discussion

In this paper, we propose an unsupervised manifold alignment algorithm, MMD-MA, for integrating multiple types of single-cell measurements carried out on disjoint populations of single cells drawn from a common source. The key advantage of our MMD-MA algorithm is that it does not require any correspondence information, either between the samples or between the features. In many real-world integration applications, such correspondence information is not available. Another advantage of our MMD-MA algorithm is that has only weak distributional requirements for the domains to be aligned, namely, that the manifolds exhibit sufficient structure to allow for alignment. This flexibility gives MMD-MA the power to potentially integrate many different types of single cell measures, including gene expression, DNA accessibility, chromatin organization, methylation, and imaging data. Furthermore, the MMD-MA framework can easily be extended to more than two domains, allowing integration of, for example, scRNA-seq, scATAC-seq, and scHi-C of single cells. We have shown that MMD-MA works well in the presence of nonlinear mappings and is robust to the choice of several hyperparameters, including the trade-off parameters, the parameters associated with the MMD term, and the dimensionality of the shared space.

Currently, MMD-MA can be used to align hundreds or thousands of single cells in a reasonable running time. The gradient descent algorithm could be parallelized to save time if multiple cores are available. For future work, we will focus on scaling up the MMD-MA algorithm. Given decreasing sequencing costs, it is likely we will need to apply MMD-MA to align millions of single cells in the future. Running MMD-MA efficiently without storing large kernel matrices in memory will be a crucial issue to solve. A possible solution may rely on random projection [12] or Nystrom approximation [16], which are approximation methods for large-scale kernel matrices.

All of the data sets used in this study, including the original and mapped simulated data and the Methyl-Expr data set, as well as the corresponding MMD-MA outputs, are available for download from http://noble.gs.washington.edu/proj/mmd-ma.

## Acknowledgments

This work was supported by National Institutes of Health award U54 DK107979.

## Supplement

**Table S1:**
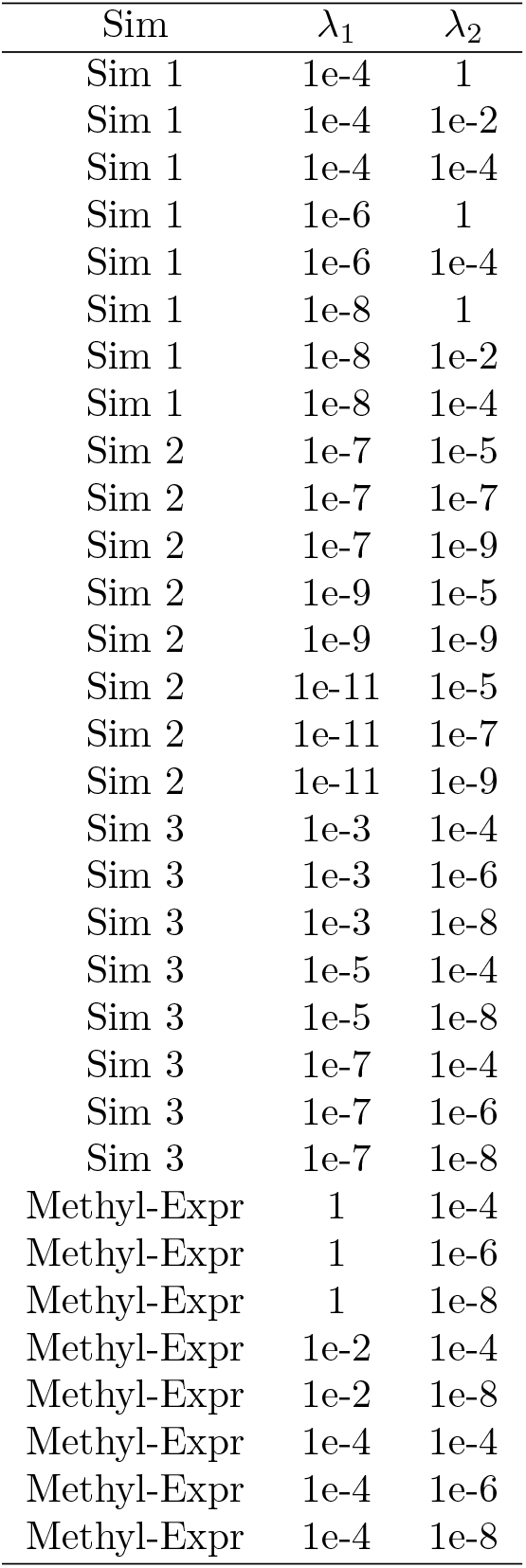
Trade-off hyperparameters investigated for each data set.

**Figure S1:**
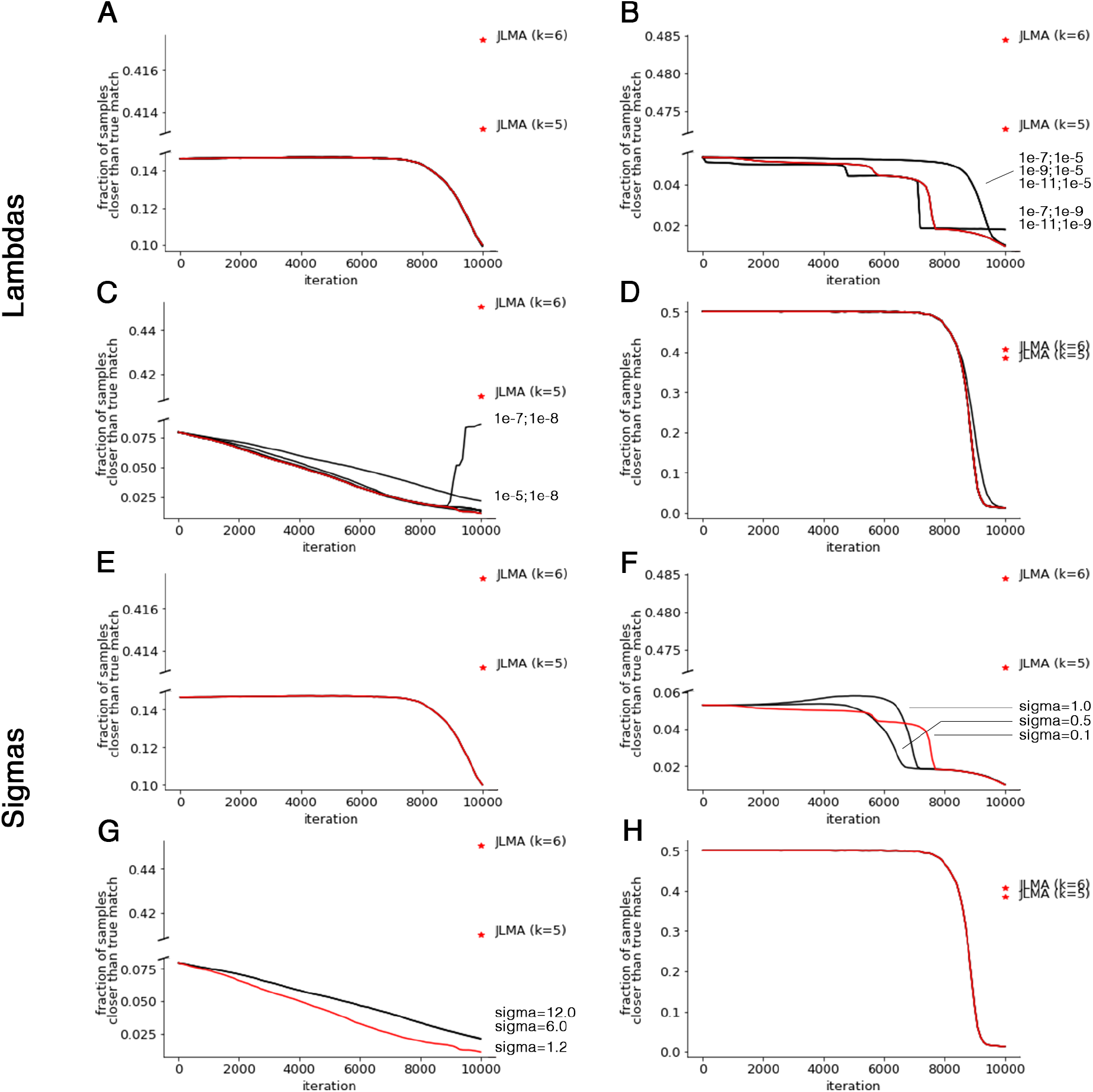
Testing sensitivity to hyperparameters. (A–D) The performance of MMD-MA when the trade-off parameters *λ*_1_ and *λ*_2_ are set differently in the three numerical simulations and the one real-world application, respectively. In each case, eight settings we chosen, and the plot only shows curves that differ from the one produced by the selected parameters. (E–H) The performance of MMD-MA when *σ* is set differently in the three numerical simulations and the one real-world application, respectively. Again, only settings that yield results different from the selected results are shown.

